# Ileal Gpbar1 gates depressive phenotypes via gut to brain “vagal-NTS-PVN” GABAergic axis

**DOI:** 10.64898/2026.09.06.749749

**Authors:** Lili Zhong, Yuanlu Shu, Min Yan, Long Wang, Min Wu, Jing Wu, Song Yang, Zedong Gu, Zhongliang Chen, Guoyan Xin, Shan Guan, Tian Tian

## Abstract

Depression represents a typical mental condition, yet whether intestinal metabolic signals drive central neural circuits remains unclear. Here we identify ileal enteric neuronal Gpbar1 as a previously unrecognized peripheral node that coordinates gut-to-brain communication to prevent depression-like behavior through a vagus-dependent, nucleus tractus solitaries (NTS)→paraventricular nucleus (PVN) signaling axis. The enteric plexus Gpbar1 is predominantly expressed on a subpopulation of GAD1-positive GABAergic neurons. Chronic restraint stress suppresses Gpbar1 expression, triggers glycolytic reprogramming and mitochondrial distress in both ileum and hypothalamus. Oral Gpbar1 agonism with INT 777 reverses hypothalamic HK2/TOM20 alterations and behavioral abnormalities by enhanced ileal mTOR/HIF-1α signaling and glycolysis-related metabolite shifts, an effect specifically abolished by subdiaphragmatic vagotomy or chemogenetic silencing of PVN GABAergic neurons. Our findings establish ileal Gpbar1 as a gut-derived metabolic sensor that engages a defined vagal–brainstem–hypothalamic inhibitory circuit to regulate depressive phenotypes, shifting the roadmap from brain-centered to gut-initiated antidepressant mechanisms and unveiling a compelling peripheral therapeutic strategy that bypasses intractable central brain-directed intervention.

## Introduction

Depression is the most prevalent psychiatric condition that affects at least 340 million people worldwide (4.4% global prevalence) and the leading cause of mental health-related disability^1, 2^, yet current pharmacotherapies—largely targeting monoaminergic systems—achieve remission in only a small fraction of patients^3^. This therapeutic bottleneck reflects an intractable dilemma: the underlying biological basis of depression is incompletely understood, and most existing treatments address downstream consequences rather than upstream pathogenic mechanisms. Chronic stress, a major contributor to depression^4, 5^, induces widespread dysfunction at the cellular, molecular and neural circuit levels^6, 7^. The heterogeneity of depressive symptomatology reflects underlying dysfunctions spanning monoaminergic neurotransmission, hypothalamic–pituitary–adrenal axis regulation, synaptic plasticity, and mitochondrial energy metabolism^8, 9, 10, 11, 12, 13^. In recent years, the gut–brain axis has emerged as a critical framework for understanding how peripheral physiological states influence central emotional processing^14, 15, 16, 17^. Pioneering studies revealed that gut microbial metabolites, such as tryptophan metabolites, short-chain fatty acids, and bile acids serve as important mediators of gut-brain communication^18, 19^. However, the precise molecular and neural mechanisms by which gut-derived signals orchestrate depression-like behaviors remain largely undefined, representing a major gap in pathophysiological understanding and a barrier to developing mechanistically informed therapies.

Among the gut–brain signaling molecules, bile acids have attracted growing attention due to their pleiotropic roles in metabolism, inflammation, and neural function^20, 21^. The G protein-coupled bile acid receptor 1 (Gpbar1, also known as TGR5) is expressed in multiple tissues, including the brain, liver, and gastrointestinal tract^22^. Central Gpbar1 has been implicated in energy homeostasis and, more recently, in emotional regulation^23, 24^. Chronic stress markedly reduces brain Gpbar1 expression through specific γ-aminobutyric acid-ergic (GABAergic) circuits involving CA3 pyramidal neurons of the mouse hippocampus^25^, whereas genetic overexpression of Gpbar1 in CA3 pyramidal neurons ameliorates depression-like phenotypes^26^. These pioneering studies established a role for central Gpbar1 in emotional regulation, yet they leave a fundamental question unanswered: Does peripheral Gpbar1 that acts as important metabolic regulators—particularly those in the enteric nervous system, a key neural component of gut-brain communication^27, 28^—contribute to stress-induced depression-like behaviors via gut-to-brain signaling?

In the present study, we address this gap by systematically investigating the role of ileal enteric neuronal Gpbar1 in a chronic restraint stress (CRS) mouse model of depression (**Supplementary Fig. 1**). We demonstrate that oral administration of the Gpbar1 agonist INT-777 robustly reverses CRS-induced anxiety- and depression-like behaviors, while concurrently normalizing hypothalamic markers of glycolysis (hexokinase 2, HK2) and mitochondrial integrity (translocase of outer membrane 20, TOM20). Using targeted energy metabolomics, we uncover coordinated glycolytic reprogramming in both the ileum and hypothalamus of CRS mice—characterized by elevated glucose-6-phosphate, dihydroxyacetone phosphate, L-lactate, and D-glucose, along with reduced acetyl-CoA—that is largely corrected by INT-777 treatment. These metabolic effects are independent of global gut microbiota remodeling, pointing instead to enteric neuronal Gpbar1 as the primary peripheral mediator.

At the cellular level, we establish that ileal Gpbar1 is predominantly localized to enteric neurons, including a GAD1-positive GABAergic subpopulation. CRS reduces Gpbar1/GAD1 colocalization and disrupts local energy metabolism within these neurons, as evidenced by increased HK2 signals and decreased TOM20 signals— abnormalities that are partially reversed by INT-777. Using pseudorabies virus (PRV)-based retrograde transsynaptic tracing from the ileal wall, we map a polysynaptic ascending pathway sequentially engaging the nodose ganglion, the nucleus tractus solitaries (NTS), and paraventricular nucleus (PVN). Remarkably, PRV-labeled neurons in both the NTS and PVN show a preferential association with GABAergic (GAD1-positive) over glutamatergic (CaMKIIα-positive) neurons, revealing a previously unrecognized ileal vagal–GABAergic axis. Functional validation by chemogenetic inhibition of PVN GABAergic neurons (via AAV-GAD67-hM4D(Gi)-mCherry) completely abolishes the antidepressant-like effects of INT-777, confirming that PVN GABAergic activity is necessary for its behavioral benefits. Furthermore, subdiaphragmatic vagotomy eliminates INT-777’s therapeutic effects, suggesting that intact vagal signaling is functionally required for gut-to-brain communication.

Collectively, our findings redefine Gpbar1 as a gut-originating metabolic gatekeeper of depression-like phenotypes, operating through an ileal enteric neuron– vagus nerve–NTS–PVN GABAergic circuit. This work provides, to our knowledge, the first demonstration that a peripheral bile acid receptor in enteric neurons modulate central emotional behavior independently of microbiota-mediated mechanisms.

## Results

### Oral Gpbar1 agonist reverses CRS-induced depression-like behaviors and corrects hypothalamic glycolysis–mitochondrial dysfunction

The CRS mouse model was successfully established after 4 weeks of chronic restraint stress (**Fig. 1A**). Compared with the naïve control (CON) group, CRS mice exhibited no significant alteration in the total distance traveled in the open field test (**Fig. 1B**), but CRS mice displayed a shorter traveling distance in the center area of the open field (**Fig. 1C**), reduced time spent in the center area (**Fig. 1D**), and a lower sucrose preference rate (**Fig. 1E**), together with prolonged immobility times in the forced swimming test and tail suspension test (**Fig. 1F,1G**). Western blot analysis further showed that Gpbar1 protein levels were significantly decreased in the hypothalamus (**Fig. 1H,1I**) of CRS mice. CRS also reduced hypothalamic PSD95 expression (**Fig. 1H,1J**) and dendritic spine density (**Fig. 1K,1L**), indicating impaired synaptic structural integrity and neuronal plasticity in the hypothalamus. In addition to the hypothalamus, CRS reduced Gpbar1 protein expression in the prefrontal cortex and hippocampus (**Supplementary Fig. 2A-D**).

**Fig. 1.**
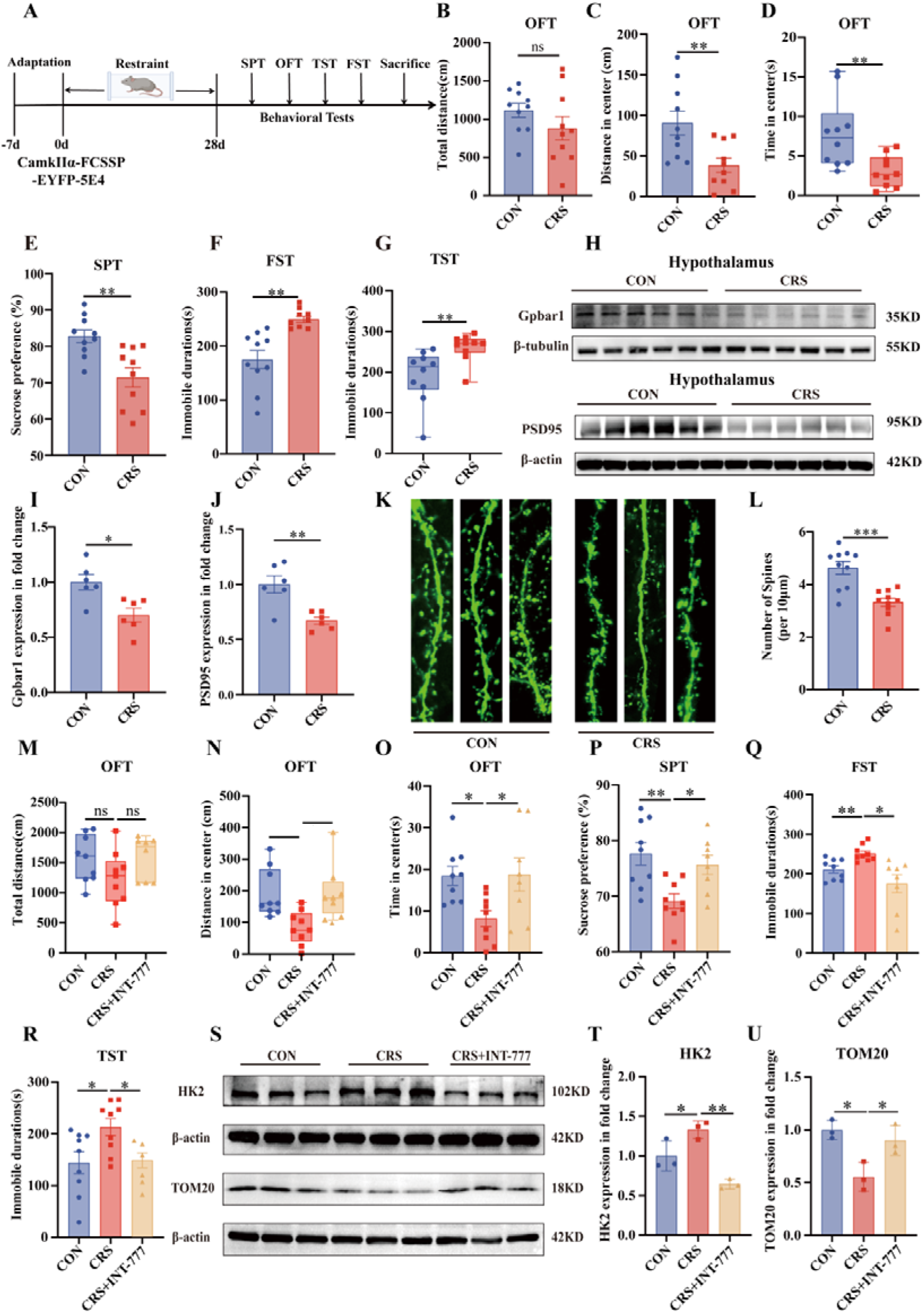
Gpbar1 activation alleviates CRS-induced depression-like behaviors and hypothalamic synaptic and metabolic abnormalities. (**A**) Experimental schematic showing viral sparse labeling (using CaMKIIα-FCSSP-EYFP-5E4 virus), chronic restraint stress (CRS) exposure, behavioral testing, and tissue collection. (**B-G**) Behavioral assessment of the naïve control (CON) and CRS mice. (**B**) Total distance traveled in the open field test (OFT) between CON and CRS mice (t = 1.135, *P* = 0.205). (**C**) Distance traveled in the center zone during the open field test (t = 3.05, *P* = 0.007). (**D**) Time spent in the center zone during the open field test (Mann–Whitney U test, *P* = 0.003). (**E**) Sucrose preference in the sucrose preference test (SPT) (t = 3.541, *P* = 0.002). (**F**) Immobility time in the forced swimming test (FST) (t = −4.213, *P* = 0.001). (**G**) Immobility time in the tail suspension test (TST) (Mann–Whitney U test, *P* = 0.001). (**H-J**) Representative Western blot images and quantification of hypothalamic Gpbar1 and PSD95 protein expression in the CON and CRS mice. (**H**) Representative immunoblots of Gpbar1 and PSD95. (**I**) Quantification of hypothalamic Gpbar1 protein levels. (**J**) Quantification of hypothalamic PSD95 protein levels. (**K-L**) Sparse-labeling analysis of hypothalamic dendritic spines. (**K**) Representative images of dendritic spines in the CON and CRS mice. (**L**) Quantification of dendritic spine density calculated as the number of spines per 10 μm dendritic segment. (**M-R**) Behavioral assessment following INT-777 administration (CRS+INT-777 group). (**M**) Total distance traveled in the OFT (H = 3.075, *P* = 0.215). (**N**) Distance traveled in the center zone during the OFT (H = 11.057, *P* = 0.004). (**O**) Time spent in the center zone during the OFT (F (2, 23) = 7.307, *P* = 0.003). (**P**) Sucrose preference in the SPT (F (2, 23) = 25.456, *P* < 0.001). (**Q**) Immobility time in the FST (F (2, 23) = 7.512, *P* = 0.003). (**R**) Immobility time in the TST (F (2, 23) = 5.285, *P* = 0.013). (**S-U**) Representative Western blots images and quantification of hypothalamic HK2 and TOM20 protein expression following INT-777 treatment. (**S**) Representative immunoblots of HK2 and TOM20. (**T**) Quantification of hypothalamic HK2 protein levels. (**U**) Quantification of hypothalamic TOM20 protein levels. For panels B-G, n = 10 mice per group. For panels M-R, sample sizes were n = 9, 9, and 8 mice per group, respectively. For Western blot analyses, n = 3-6 mice per group as indicated in the figure. For dendritic spine analysis, n = 3 mice per group, and 10 dendritic segments were analyzed in each group. ns, not significant; \**P* < 0.05, \*\**P* < 0.01, and \*\*\**P* < 0.001.

To assess the effect of Gpbar1 activation, mice received oral gavage of a Gpbar1 agonist INT-777 for the final 14 days of the 4-week CRS procedure, followed by behavioral testing after CRS completion (**Supplementary Fig. 2E**). No significant difference in total distance traveled in the open field was observed among the three groups (**Fig. 1M**). Compared with the CON group, CRS mice showed reduced distance traveled and time spent in the center area, decreased sucrose preference, and increased immobility times in the forced swimming and tail suspension tests (**Fig. 1N-R**), all of which were significantly reversed by INT-777 treatment. Consistently, locomotor heat maps depict blunted center-area exploration in the CRS cohort, whereas pharmacological Gpbar1 activation by INT-777 robustly elevates center-zone activity (**Supplementary Fig. 2F**). Body weight changed similarly among the three groups during the experimental period, indicating that INT-777 is safe and biocompatible (**Supplementary Fig. 2G**).

Previous studies suggest that Gpbar1 activation can increase energy expenditure and improve glucose tolerance ^29, 30^. The hypothalamus is a key brain region involved in the regulation of food intake, energy expenditure, and glucose metabolism, and is therefore highly relevant to central metabolic alterations^30, 31^. We further examined hypothalamic HK2 and TOM20 protein expression to assess the effects of Gpbar1 activation after CRS on glycolysis- and mitochondria-related indices. Results show that CRS stimulation increased HK2 expression while reducing TOM20 levels in the hypothalamus (**Fig. 1S-U**), suggesting enhanced glycolytic metabolism accompanied by mitochondrial dysfunction. Notably, INT-777 treatment decreased HK2 expression and restored TOM20 abundance in CRS mice, indicating that Gpbar1 activation mitigates chronic restraint stress-induced metabolic reprogramming and improves mitochondrial homeostasis.

### Ileal Gpbar1 activation alleviates depression via gut–brain metabolic crosstalk independent of microbiota remodeling

Gpbar1 is expressed throughout the intestine, particularly in the ileum, colon, and enteric nervous system ^32, 33^. Since the ileum is a major intestinal segment for bile acid reabsorption and active bile acid receptor signaling^34^. We first examined whether INT-777 mediated Gpbar1 activation affected CRS-induced alterations in the ileal microenvironment. 16S rRNA sequencing revealed that CRS significantly reduced ileal microbial α-diversity, as indicated by decreased Chao, Shannon, and Simpson indices compared with CON mice (**Supplementary Fig. 3A-C**). Surprisingly, INT-777 treatment fails to restore these metrics (**Supplementary Fig. 3A-C**), indicating that the compound cannot reverse the CRS-induced loss of ileal microbiota richness and diversity. Principal coordinates analysis (PCoA) showed a separation trend among the three groups, whereas the Adonis test did not reach statistical significance (R² = 0.16, P = 0.054; **Supplementary Fig. 3D**), suggesting that INT-777 did not induce a significant overall change in microbial beta diversity. Further taxonomic analysis showed that CRS altered the composition of the ileal microbiota. At the phylum level, the relative abundance of several taxa showed a trend toward normalization after INT-777 treatment, indicating a modest regulatory effect on higher-level microbial composition (**Fig. 2A**). However, at the genus level, the CRS+INT-777 group did not show broad or consistent microbial reconstruction and remained overall closer to the CRS group (**Fig. 2B**). Taken together, these results indicate that CRS induced compositional disruption and diminished diversity of the ileal microbiota. Although INT-777 exerted a limited modulatory effect at the phylum level, it neither significantly restore alpha or beta diversity, nor did it produce broad changes at the genus level, its therapeutic effects are unlikely to rely primarily on global remodeling of the ileal microbial structure.

**Fig. 2.**
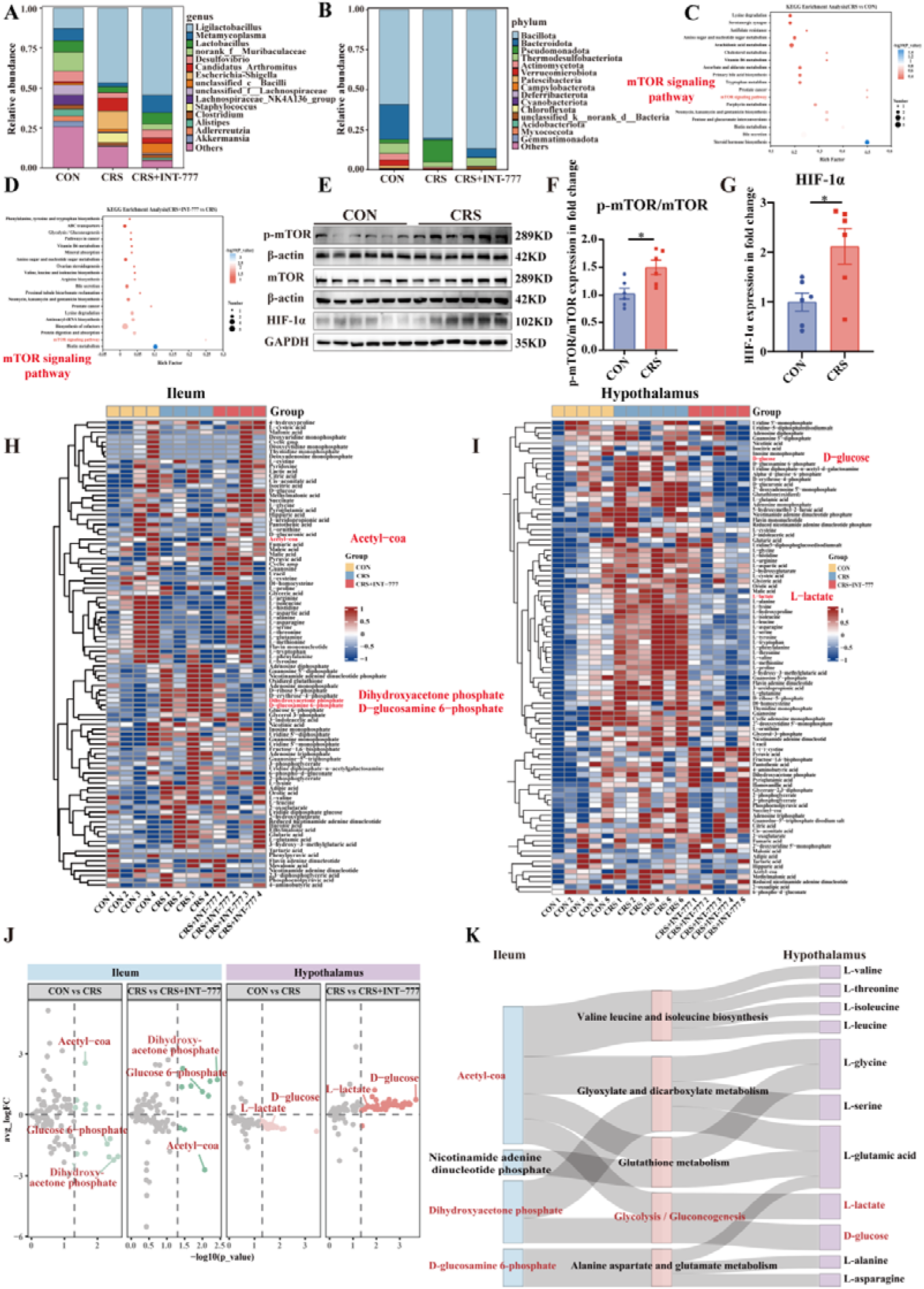
Gpbar1 activation modulates ileal microbiota-associated mTOR/HIF-1 α signaling and ileum–hypothalamus energy metabolism in CRS mice. (**A**) Relative abundance of the dominant bacterial genera in ileal samples from CON, CRS, and CRS+INT-777 mice. (**B**) Relative abundance of the dominant bacterial phyla in ileal samples from CON, CRS, and CRS+INT-777 mice. (**C**) KEGG pathway enrichment analysis of differential ileal metabolites between CRS and CON mice. (**D**) KEGG pathway enrichment analysis of differential ileal metabolites between CRS+INT-777 and CRS mice. The mTOR signaling pathway was enriched in both comparisons. (**E**) Representative Western blots of p-mTOR, total mTOR, and HIF-1α in ileal tissues from CON and CRS mice. (**F**) Quantification of p-mTOR expression in ileal tissues. (**G**) Quantification of HIF-1α expression in ileal tissues. (**H**) Hierarchical clustering heatmap of targeted energy metabolites in ileal samples from CON, CRS, and CRS+INT-777 mice. (**I**) Hierarchical clustering heatmap of targeted energy metabolites in hypothalamic samples from CON, CRS, and CRS+INT-777 mice. (**J**) Differential distribution of targeted energy metabolites in the ileum and hypothalamus across pairwise group comparisons. Representative altered metabolites, including glucose 6-phosphate, dihydroxyacetone phosphate, acetyl-CoA, L-lactate, and D-glucose, are labeled. (**K**) Sankey diagram illustrating metabolic network connectivity between ileal and hypothalamic differential metabolites. The left column represents differential metabolites in the ileum (blue), the middle column represents enriched metabolic pathways (pink), and the right column represents differential metabolites in the hypothalamus (purple). The width of connecting bands represents the relative contribution of each metabolic association. Sample sizes were n = 4, 4, and 4 mice per group for ileal metabolomics and n = 5, 6, and 5 mice per group for hypothalamic targeted energy metabolomics. ns, not significant; \**P* < 0.05, \*\**P* < 0.01, and \*\*\**P* < 0.001.

We therefore identified whether INT-777 regulates ileal metabolic states independent of microbial reconstruction. Untargeted metabolomics identified increased ileal L-leucine levels in CRS mice, which were reduced following INT-777 treatment (**Supplementary Fig. 3E-F**). The Kyoto Encyclopedia of Genes and Genomes (KEGG) enrichment analysis of differential metabolites revealed enrichment of the mTOR signaling pathway in both CRS versus CON and CRS+INT-777 versus CRS comparisons (**Fig. 2C,2D**). Consistent with this pathway-level finding, CRS increased ileal p-mTOR/mTOR ratios and HIF-1α expression compared with CON mice (**Fig. 2E–G**).

Targeted energy metabolomics revealed distinct metabolic profiles in ileal and hypothalamic tissues among the three groups (**Fig. 2H,2I**). A total of 96 and 93 differential energy metabolites were identified in the ileum and hypothalamus, respectively. CRS increased glucose-6-phosphate and dihydroxyacetone phosphate while reducing acetyl-CoA in the ileum (**Fig. 2H**). Meanwhile, D-glucose and L-lactate were elevated in the hypothalamus (**Fig. 2I**). These metabolic changes were reversed after INT-777 treatment (**Fig. 2J**). Sankey network analysis showed metabolic network connectivity between ileal and hypothalamic metabolites were mainly enriched in glycolysis/gluconeogenesis, glyoxylate and dicarboxylate metabolism, glutathione metabolism, branched-chain amino acid biosynthesis, and amino acid metabolism pathways (**Fig. 2K**). These findings indicate that INT-777 modulates coordinated metabolic alterations in the ileum and hypothalamus without broadly restoring the ileal microbiota.

### Enteric Gpbar1 resides in a GAD1-positive GABAergic subpopulation susceptible to stress-induced disruption

We next examined the cellular localization of Gpbar1 in the ileal enteric nervous system and its relationship with critical enteric neurons. Immunofluorescence co-staining of Gpbar1 with the enteric neuronal markers PGP9.5 and TUJ1 revealed that, in the myenteric plexus (MP), Gpbar1 signals were mainly distributed in PGP9.5- and TUJ1-positive neuronal structures, with limited colocalization with the enteric glial marker S100β (**Fig. 3A**). Quantification using the Manders colocalization coefficient confirmed this observation (**Fig. 3B**). A similar trend was observed in the submucosal plexus (SMP) (**Fig. 3C,3D**). These findings suggest that Gpbar1 is primarily localized to ileal enteric neurons rather than enteric glial cells.

**Fig. 3.**
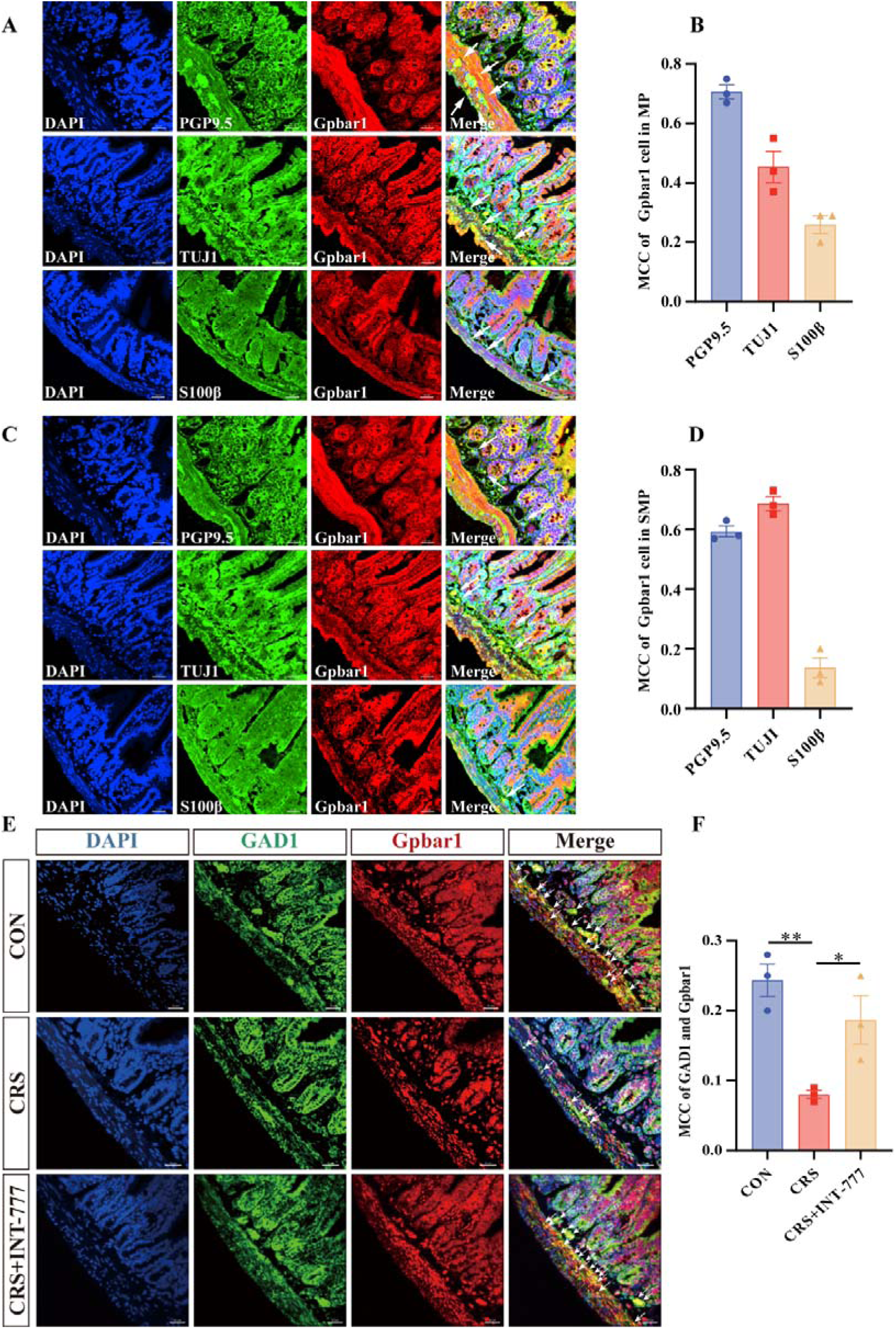
Enteric neuronal localization of Gpbar1 in the ileal plexuses and modulation of depression-like behavior by GABAergic Gpbar1. (**A**) Representative immunofluorescence images showing colocalization of Gpbar1 with S100β, TUJ1, and PGP9.5 in the ileal enteric nervous system. DAPI, blue; S100β, TUJ1, or PGP9.5, green; Gpbar1, red. Scale bar = 30 μm. (**B**) Quantification of Manders colocalization coefficients for Gpbar1 with PGP9.5, TUJ1, and S100β in the myenteric plexus. (**C**) Representative immunofluorescence images showing colocalization of Gpbar1 with PGP9.5, TUJ1, and S100β in the submucosal plexus. (**D**) Quantification of Manders colocalization coefficients for Gpbar1 with PGP9.5, TUJ1, and S100β in the submucosal plexus. (**E**) Representative immunofluorescence images showing colocalization of Gpbar1 with GAD1 in ileal tissues from CON, CRS, and CRS+INT-777 mice. DAPI, blue; GAD1, green; Gpbar1, red. Scale bar = 30 μm. (**F**) Quantification of Manders colocalization coefficients between Gpbar1 and GAD1 in ileal tissues. n = 3 mice per group. ns, not significant; \**P* < 0.05, \*\**P* < 0.01, and \*\*\**P* < 0.001.

Previous studies have shown that GABA (γ-aminobutyric acid)-positive neurons can be detected in the myenteric plexus of the mouse ileum, indicating that ileal GABAergic neurons represent a neurochemical subpopulation within the enteric nervous system^35, 36^. GAD1/GAD67, key enzymes involved in GABA synthesis^37^, are commonly used as markers of GABAergic neurons ^36^. We then investigated the relationship between Gpbar1 and GAD1-positive enteric neurons. GAD1/Gpbar1 co-staining showed that a subset of Gpbar1-positive signals was located within GAD1-positive enteric neuronal structures, indicating that Gpbar1-positive enteric neurons in the ileum include a GAD1-positive GABAergic subpopulation (Fig. 3E). In contrast, Gpbar1/GAD1 colocalization was reduced in the CRS group compared to the CON group, whereas INT-777 treatment increased Gpbar1/GAD1 colocalization to a similar level that of CON group (**Fig. 3E,3F**). These results suggest that CRS disrupts Gpbar1-related signaling in GAD1-positive enteric neurons in the ileum, with Gpbar1 activation by INT-777 can partially relieve this alteration.

### Gpbar1 activation reverses glycolytic–mitochondrial metabolic reprogramming in ileal enteric neurons

To investigate whether CRS disrupts Gpbar1-associated metabolic homeostasis in enteric neurons and whether Gpbar1 activation restores neuronal metabolic balance, we assessed glycolytic and mitochondrial alterations in Gpbar1-positive enteric neuronal structures. Gpbar1 signals in PGP9.5-positive enteric neuronal structures were decreased in ileal tissue from the CRS group compared with the CON group and were increased after INT-777 intervention (**Fig. 4A-C**). Hexokinase 2 (HK2), which catalyzes the first step of glycolysis and regulates glycolytic flux ^38^, was increased in PGP9.5-positive regions after CRS and reduced by INT-777 (**Fig. 4A,4D**). ELISA further showed that lactate and HK2 levels in ileal tissue were increased in the CRS group and decreased after INT-777 intervention (**Supplementary Fig. 4A,4B**). By comparison, translocase of outer mitochondrial membrane 20 (TOM20), a receptor component of the mitochondrial protein import machinery^39^, was reduced in PGP9.5-positive regions after CRS and restored by INT-777 (**Fig. 4B,4F**). Taken together, INT-777 attenuated glycolytic remodeling in the CRS group while restoring mitochondrial signatures in ileal enteric neurons.

**Fig. 4.**
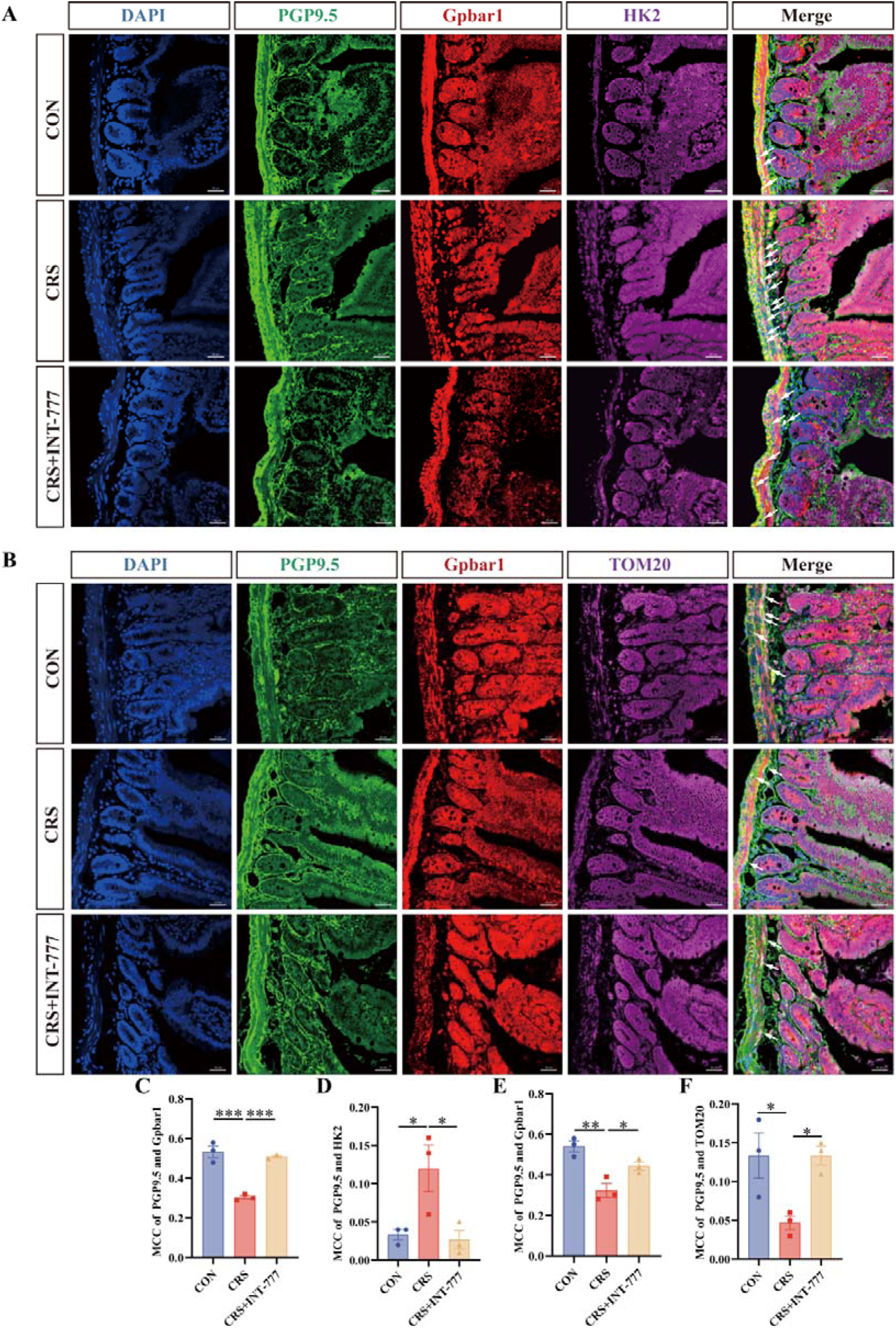
INT-777 attenuated glycolytic remodeling while restoring mitochondrial signatures in ileal enteric neurons. (**A**) Representative immunofluorescence images of DAPI (blue), PGP9.5 (green), Gpbar1 (red), HK2 (purple), and merged signals in ileal tissues from the three groups. Scale bar = 25 μm. (**B**) Representative immunofluorescence images of DAPI (blue), PGP9.5 (green), Gpbar1 (red), TOM20 (purple), and merged signals in ileal tissues from the three groups. Scale bar = 25 μm. (**C**) Quantification of Gpbar1 signals within PGP9.5-positive regions in the ileum. Gpbar1 signals were decreased in the CRS group compared with the CON group and increased after INT-777 oral gavage. (**D**) Quantification of HK2 signals within PGP9.5-positive regions in the ileum. HK2 signals were increased in the CRS group compared with the CON group and decreased after INT-777 oral gavage. (**E**) Quantification of Gpbar1 signals within PGP9.5-positive regions in the ileum. Gpbar1 signals were decreased in the CRS group compared with the CON group and increased after INT-777 oral gavage. (**F**) Quantification of TOM20 signals within PGP9.5-positive regions in the ileum. TOM20 signals were decreased in the CRS group compared with the CON group and increased after INT-777 gavage. n = 3 mice per group. ns, not significant; \**P* < 0.05, \*\**P* < 0.01, and \*\*\**P* < 0.001.

### Ileal PRV tracing reveals a gut–brain GABAergic circuit from ileal neurons to PVN

To trace the neuroanatomical connections from the ileum to the central nervous system, we injected PRV-CAG-EGFP, a neurotropic pseudorabies virus used for retrograde transsynaptic tracing, into the ileal wall (**Fig. 5A**). Enhanced green fluorescent protein positive (EGFP^+^) signals were detected in local neural structures of the ileum (**Fig. 5B,5C**), and subsequently in the nodose ganglion (NG), nucleus tractus solitarius (NTS), and paraventricular nucleus of the hypothalamus (PVN) (**Fig. 5D-F**), indicating the presence of polysynaptic connections involve the NG, NTS, and PVN. This anatomical distribution is in line with previous herpes simplex virus (HSV)-based anterograde transsynaptic tracing showing that gut vagal sensory signals can access NTS GABAergic neurons projecting to the PVN^40^.

**Fig. 5.**
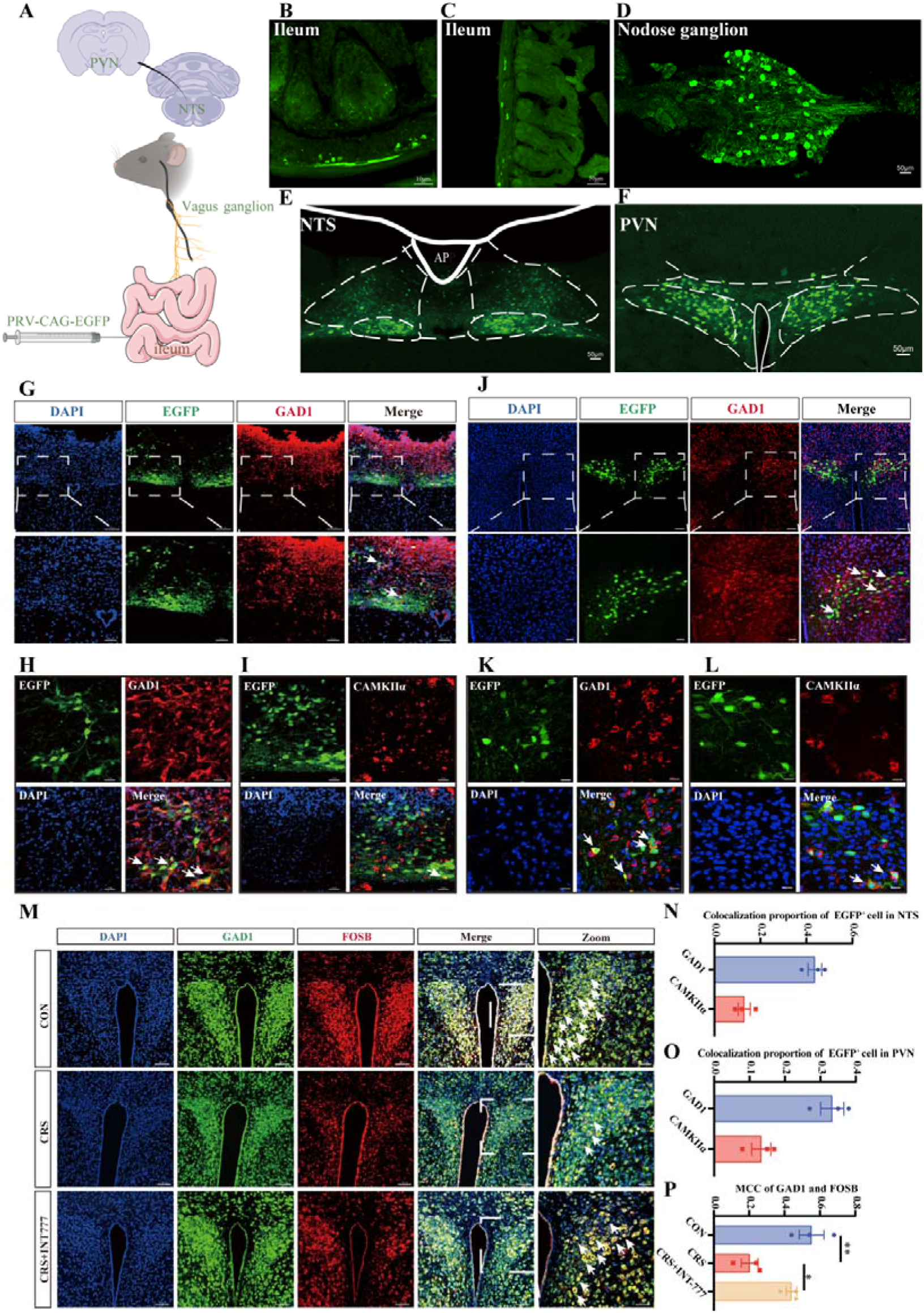
Gut-derived PRV signals are preferentially associated with NTS and PVN GABAergic neurons in the ileum–brain pathway. (**A**) Schematic diagram of PRV-CAG-EGFP (a neurotropic pseudorabies virus expressing enhanced green fluorescent protein)-mediated retrograde transsynaptic tracing from the ileum to the PVN. (**B–C**) Representative images of EGFP^+^ signals in the ileum. (**D**) Representative image of EGFP^+^ signals in the nodose ganglion. (**E**) Representative image of EGFP^+^ signals in the NTS. (**F**) Representative image of EGFP-positive signals in the PVN. n = 3 mice. (**G**) Co-staining of GAD1 and PRV-EGFP-positive signals in the NTS at 20× and 40× magnification. Scale bars = 100 μm and 50 μm, respectively. (**H**) Co-staining of GAD1 and PRV-EGFP^+^ signals in the NTS. Scale bar = 25 μm. (**I**) Co-staining of CaMKIIα and PRV-EGFP^+^ signals in the NTS. Scale bar = 25 μm. (**J**) Co-staining of GAD1 and PRV-EGFP^+^ signals in the PVN at 20× and 40× magnification. Scale bars = 100 μm and 50 μm, respectively. (**K**) Co-staining of GAD1 and PRV-EGFP^+^ signals in the PVN. Scale bar = 25 μm. (**L**) Co-staining of CaMKIIα and PRV-EGFP^+^ signals in the PVN. Scale bar = 25 μm. (**M**) Representative immunofluorescence images of GAD1 and FosB in the PVN across the three groups. Scale bar = 100 μm; zoomed images, scale bar = 50 μm. n = 3 mice per group. (**N**) Quantification of EGFP^+^ signal co-localization with GAD1 and CaMKIIα in the NTS. n = 3 mice. (**O**) Quantification of EGFP^+^ signal co-localization with GAD1 and CaMKIIα in the PVN. n = 3 mice. (**P**) Quantification of FosB signals within GAD1^+^ regions in the PVN. FosB signals within GAD1^+^ regions were decreased in the CRS group compared with the CON group and increased after INT-777 oral gavage. ns, not significant; \**P* < 0.05, \*\**P* < 0.01, and \*\*\**P* < 0.001.

We further characterized the neurochemical phenotype of tracer-labeled cells by co-staining EGFP^+^ cells with excitatory neuronal markers CamKIIα and the inhibitory neuronal marker GAD1. In the NTS, EGFP^+^ signals showed greater colocalization with GAD1-positive cells than with CaMKIIα-positive cells (**Fig. 5G–I**). In the PVN, EGFP^+^ signals also displayed superior colocalization with GAD1-positive cells compared to CaMKIIα-positive counterpart (**Fig. 5J–L**). Quantification confirmed greater colocalization of EGFP+ signals with GAD1 than with CaMKIIα in both the NTS and PVN (**Fig. 5N,5O)**, suggesting that PRV-labeled neurons in the NTS and PVN were preferentially associated with GABAergic neuronal populations.

Inspired by the above PRV tracing results, we further analyzed FosB signals in the PVN and their relationship with GAD1 or CaMKIIα. FosB is a classical immediate early gene^41^. And its relatively long half-life and sustained stable expression make it suitable for evaluating neuronal activation following chronic stimulation^42^. Immunofluorescence analysis in the PVN revealed more frequent colocalization of FosB with the GABAergic neuronal marker GAD1 and less colocalization with CaMKIIα (**Supplementary Fig. 5A,5B**). GAD1/FosB co-staining further showed reduced colocalization in CRS mice compared with CON mice, whereas INT-777 treatment restored this signal (**Fig. 5M,5P**), indicating that INT-777 oral gavage improved the activation of GABAergic neurons.

### Functional validation of the ileal–vagal–PVN GABAergic axis in antidepressant-like effects

Considering both PRV tracing and FosB staining pointed to PVN GABAergic neurons, we therefore adopted the inhibitory hM4D(Gi) designer receptor, a Gi-coupled chemogenetic receptor that suppresses neuronal activity exclusively upon agonist activation by a designer drug (DREADD), for loss-of-function validation^43^. The experimental design is shown in **Fig. 6A**. Wild-type C57BL/6J mice received bilateral stereotaxic injection of AAV-GAD67-hM4D(Gi)-mCherry into the PVN (hM4D(Gi) group), whereas control mice received AAV-GAD67-mCherry into the same region (CON group) (**Fig. 6B**). After recovery of viral expression, mice assigned to the CRS groups underwent CRS modeling, and INT-777 was administered to the CRS+INT-777 and CRS+INT-777+hM4D(Gi) groups. Deschloroclozapine (DCZ), a potent DREADD agonist, was administered before behavioral testing to inhibit hM4D(Gi)-expressing PVN GABAergic neurons^44^. Immunofluorescence showed that mCherry signals were expressed in the PVN and colocalized with GAD1-positive neurons, confirming viral expression in PVN GABAergic neurons (**Fig. 6C,6D**). In the open field test, no significant difference in total distance traveled was observed among groups, indicating no marked overall locomotor deficit (**Fig. 6E**). Compared with the CRS group, INT-777 intervention increased the distance traveled and time spent in the center area, whereas these effects were markedly attenuated after hM4D(Gi)-mediated inhibition of PVN GABAergic neurons (**Fig. 6F,6G**). Although INT-777 increased sucrose preference in CRS mice, this effect was abolished after inhibition of PVN GABAergic neurons (**Fig. 6H**). In the forced swimming and tail suspension tests, immobility time was increased in the CRS group and decreased after INT-777 intervention, whereas hM4D(Gi)-mediated inhibition attenuated the effect of INT-777 on reducing immobility time (**Fig. 6I,6J**). These results suggest that the activity of PVN GABAergic neurons is required for the antidepressant-like effect of INT-777 in CRS mice.

**Fig. 6.**
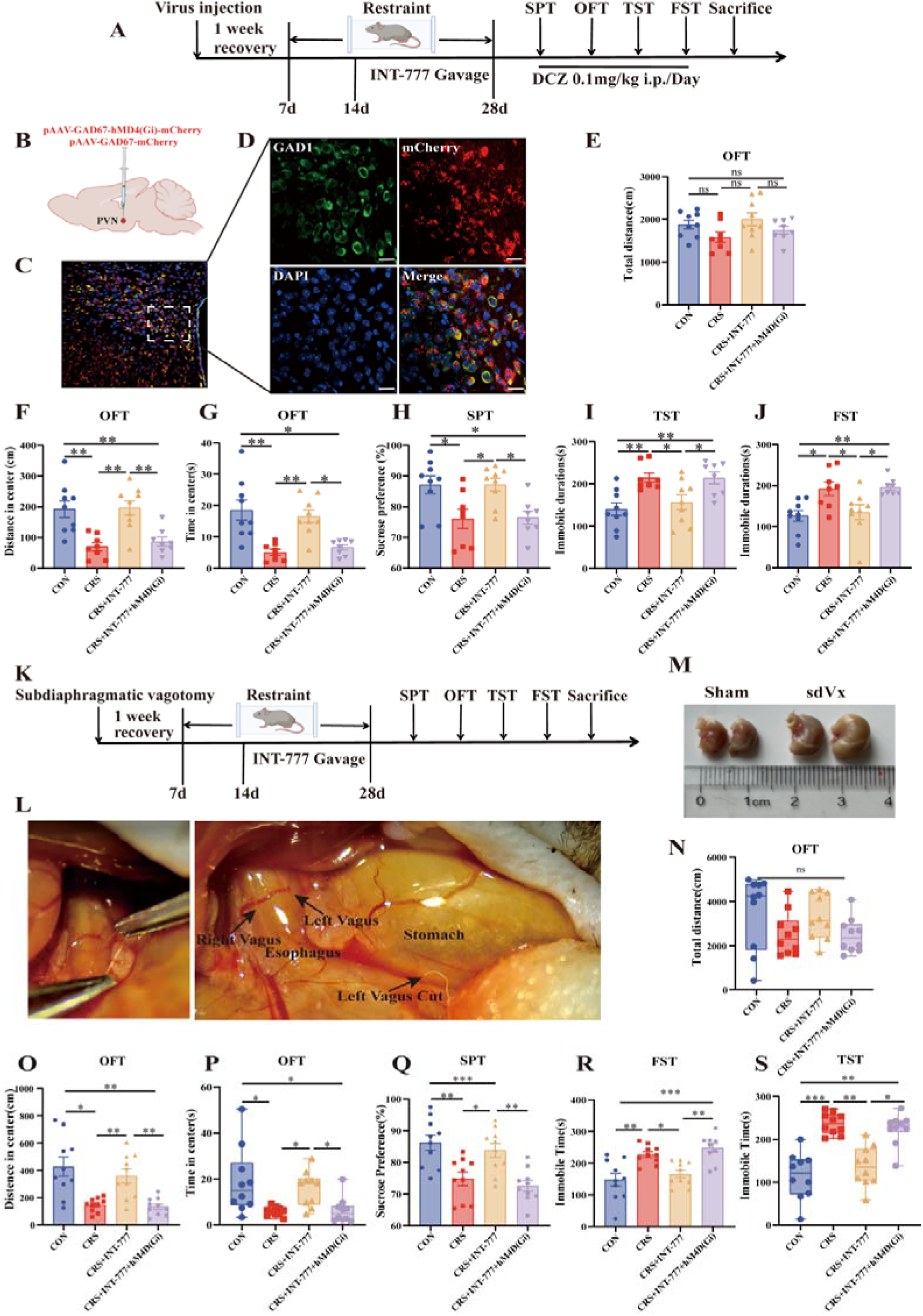
Chemogenetic inhibition of PVN GABAergic neurons and subdiaphragmatic vagotomy attenuate the antidepressant-like effects of Gpbar1 activation in CRS mice. (**A**) Timeline of viral injection, CRS modeling, INT-777 gavage, and behavioral testing. (**B**) Schematic diagram of viral injection into the PVN. (**C**) Co-localization of viral mCherry fluorescence with GAD1 in the PVN. Scale bar=50 μm. (**D**) Co-localization of viral mCherry fluorescence with GAD1 in the PVN. DAPI, blue; GAD1, green; mCherry, red. Scale bar=10 μm. (**E**) Total distance traveled in the OFT (F(3,30)=2.268, *P*>0.05). (**F**) Distance traveled in the center zone during the OFT (F(3,30)=10.19, *P*<0.001). (**G**) Time spent in the center zone during the OFT (F(3,30) = 8.764, *P*=0.002). (**H**) Sucrose preference in the SPT (F(3,30) = 5.883, *P*=0.002). (**I**) Immobility time in the FST (F(3,30) = 6.434, *P*=0.001). (**J**) Immobility time in the TST (F(3,30) = 6.905, *P*=0.001). Sample sizes were n = 9, 8, 9, and 8 mice per group. (**K**) Timeline of subdiaphragmatic vagotomy (sdVx), CRS modeling, INT-777 oral gavage, and behavioral testing. (**L**) Representative surgical images showing exposure and transection of the subdiaphragmatic vagal trunks. (**M**) Representative stomach images from sham-operated and sdVx mice. (**N**) Total distance traveled in the OFT. (**O**) Distance traveled in the center zone during the OFT. (**P**) Time spent in the center zone during the OFT. (**Q**) Sucrose preference in the SPT. (**R**) Immobility time in the FST. (**S**) Immobility time in the TST. Sample sizes were n = 10, 10, 10, and 10 mice per group. ns, not significant; \**P* < 0.05, \*\**P* < 0.01, and \*\*\**P* < 0.001.

To determine whether the vagus nerve is involved in the INT-777-mediated gut-brain axis, we performed subdiaphragmatic vagotomy (sdVx) before CRS modeling and INT-777 treatment (**Fig. 6K**). The subdiaphragmatic vagal trunks were fully exposed and transected during surgery (**Fig. 6L**). Compared with sham-operated mice, sdVx mice showed marked gastric distension and retention of gastric contents after surgery, consistent with impaired gastric emptying following subdiaphragmatic vagotomy (**Fig. 6M**). Behavioral changes in the CON, CRS, and CRS+INT-777 groups were consistent with those observed in the preceding experiment (**Fig. 6N–S**). Notably, compared with the CRS+INT-777 group, the CRS+INT-777+sdVx group showed reduced distance traveled and time spent in the center area, decreased sucrose preference, and increased immobility times in the forced swimming and tail suspension tests while total distance traveled in the open field test remained unchanged (**Fig. 6N–S**), indicating that sdVx attenuated the behavioral effects of INT-777 and supporting the involvement of the vagus nerve in INT-777-mediated gut-brain signaling.

## Discussion

This study identifies ileal enteric neuronal Gpbar1 and a vagus nerve-dependent pathway involving GABAergic neurons in the nucleus tractus solitarius (NTS) and paraventricular nucleus of the hypothalamus (PVN) that regulates depression-like behaviors. This finding extends the functional scope of Gpbar1 from previously described central mood-related circuits toward peripheral enteric metabolic sensing. Previous studies have mainly implicated Gpbar1 in hippocampal and hypothalamic neurons^25, 35, 36^. Our findings add a critical peripheral component to this framework, suggesting that enteric neuronal Gpbar1 may engage vagal interoceptive signaling in parallel with central Gpbar1 pathways.

Multi-omics data further suggest that depression is not restricted to a single brain region. It is often accompanied by coordinated changes in gut metabolite profiles, brain function, and emotional phenotypes ^15, 45^. In our study, CRS altered the diversity and composition of the gut microbiota. However, the Gpbar1 agonist INT-777 did not restore the microbial structure in CRS mice. This suggests that its antidepressant-like effect is probably independent of reshaping the gut microbiota. Previous gut-brain axis studies have emphasized microbial changes as an important source of peripheral signals. Yet the metabolic state of intestinal tissue, enteric neuronal activity, and vagal afferent input can also relay peripheral information to the brain^46, 47^. We therefore did not attribute the effect of INT-777 to microbial changes, but instead pinpointed ileal metabolite shifts and enteric neuronal Gpbar1 signaling as its primary functional targets. Our findings suggest that activation of ileal Gpbar1 may contribute to antidepressant-like effects through metabolic remodeling along the ileum– hypothalamus axis and vagus-dependent signaling, rather than through restoration of gut microbial composition.

We next linked this metabolic phenotype to vagal neural transmission. The vagus nerve provides a major route by which visceral information reaches the NTS, which subsequently engages hypothalamic networks involved in homeostatic and stress-related regulation^48, 49, 50, 51^. Our PRV tracing identified an ileum-associated polysynaptic network involving the nodose ganglion, NTS and PVN, with preferential association of tracer-labeled neurons with GABAergic populations. Importantly, recent anterograde tracing independently demonstrated that upper-gut vagal sensory neurons directly recruit NTS GABAergic neurons and that NTS GABAergic projections reach the PVH/PVN^40^. Together with the loss of INT-777 efficacy following subdiaphragmatic vagotomy and chemogenetic inhibition of PVN GABAergic neurons, these observations support a vagus-dependent GABAergic component linking peripheral Gpbar1 signaling to central stress-related circuitry and suggest that the behavioral effect may depend on the functional organization of specific inhibitory circuits rather than on a generalized change in GABAergic tone. An apparent paradox arises from the requirement for GABAergic neuronal activity in our model. Li et al. showed that chronic stress increases the excitability of GABAergic neurons in the lateral hypothalamic area (LHA) and that Gpbar1 activation alleviates depressive-like behavior by suppressing this hyperexcitability within the LHA^GABA^–dorsal CA3 (dCA3)^CaMKIIα^–dorsolateral septum (DLS)^GABA^ circuit^25^. In contrast, our results revealed the opposite directional change in PVN-associated GABAergic activity: CRS reduced FosB/GAD1 colocalization, whereas INT-777 restored it, and chemogenetic inhibition of PVN GABAergic neurons abolished the behavioral benefit. Rather than indicating contradictory roles of GABA, these findings are more consistent with the circuit- and neuronal population-specific organization of inhibitory signaling, in which the behavioral consequence of GABAergic activity depends on the identity of the downstream target and the functional position of inhibitory neurons within a given circuit^52^. Directionally divergent GABAergic effects have also been observed across depression-related circuits. For example, potentiation of basolateral amygdala GABAergic neurons suppresses excitatory basolateral amygdala (BLA)^CaMKIIα^ neurons and alleviates stress-induced depressive-like behavior^53^, whereas hyperexcitability of LHA GABAergic neurons promotes depressive-like behavior in the Gpbar1-related LHA circuit^25^. Consistent with this circuit dependence, excitation–inhibition abnormalities in depression are regionally heterogeneous, with GABAergic signaling showing different directional changes across brain regions and networks^54^. In the PVN, corticotropin-releasing hormone (CRH) neurons constitute a major stress-effector population controlling hypothalamic–pituitary–adrenal axis output, and chronic stress can shift their synaptic balance toward excitation by increasing excitatory drive and weakening GABAergic inhibitory control^55^. Moreover, recent circuit evidence demonstrates that peri-PVN GABAergic corticotropin-releasing factor receptor 1 (CRFR1) neurons can suppress PVN^CRF^ activity and mitigate stress-related behavioral responses^56^. Taken together, these observations argue against interpreting Gpbar1-related antidepressant effects as requiring a uniform increase or decrease in GABAergic activity. Instead, Gpbar1-related signaling may normalize circuit-specific inhibitory states: suppressing pathological GABAergic hyperexcitability in the LHA in the previous study^25^, while restoring insufficient inhibitory restraint within PVN-associated stress circuitry in our model. Thus, the apparently opposite changes in LHA and PVN GABAergic activity may represent different forms of circuit-specific correction of stress-induced excitation–inhibition imbalance rather than opposing actions of GABA itself. The recent observation that norepinephrine reuptake inhibition can increase tonic norepinephrine while suppressing stress-evoked phasic norepinephrine in the lateral habenula provides a useful conceptual precedent: the behavioral effect of a neurotransmitter depends on circuit context and activity dynamics, rather than its overall level alone^57^. However, because GABA release dynamics were not directly measured here, tonic–phasic regulation should not be inferred from our data.

Taken together, our findings support a peripheral metabolic–neural component of Gpbar1-mediated behavioral regulation. The present findings position enteric Gpbar1 at the interface between intestinal metabolism, vagal interoception and central stress-related inhibitory circuitry, providing a framework for investigating peripheral bile acid receptor signaling in depressive disorders.

### Conclusion

In summary, the present study indicates that pharmacological activation of Gpbar1 in the ileum of enteric plexus by INT-777 alleviates depression-like behaviors in CRS mice, improves glycolysis-related metabolic abnormalities and ameliorates energy metabolic abnormalities in both the ileum and hypothalamus. INT-777-mediated Gpbar1 activation reduced HK2 and lactate levels in ileal tissue from CRS mice, while decreasing HK2 signals and restoring TOM20 signals in Gpbar1-positive enteric neuronal structures. The behavioral benefits of Gpbar1 activation by INT-777 are depend on the integrity of subdiaphragmatic vagal signaling and on GABAergic neurons activity in the PVN, demonstrating the role of an ileal vagal afferent pathway linking the NTS with PVN GABAergic neurons. Overall, this study deepens our understanding of peripheral Gpbar1 activation and identifies PVN GABAergic neurons in gut as potential therapeutic targets for antidepressant strategies.

## Materials and methods

### Ethics statement

All animal procedures were approved by the Animal Ethics Committee of Guizhou Medical University (approval number: 2000758) and performed in accordance with the ARRIVE 2.0 guidelines. Efforts were made to minimize animal suffering and reduce the number of animals used.

### Animals

Male wild type C57BL/6J mice aged 8-12 weeks and weighing 18-25 g were obtained from the Animal Experimental Center of Guizhou Medical University. Mice were housed individually under controlled conditions with a 12 h light/dark cycle, temperature of 22-25 , and free access to food and water. Animals were acclimated for 1 week before experiments.

### Experimental design and grouping

Mice were randomly assigned to different experimental cohorts according to the experimental purpose. Separate cohorts were used for CRS model validation, INT-777 gavage, PRV-mediated tracing, PVN GABAergic chemogenetic inhibition, and subdiaphragmatic vagotomy. For CRS model validation, mice were assigned to CON and CRS groups. For INT-777 gavage, mice were assigned to CON, CRS, and CRS + INT-777 groups. For chemogenetic experiments, mice were assigned to CON, CRS, CRS + INT-777, and CRS + INT-777 + hM4D(Gi) groups. For vagotomy experiments, mice were assigned to CON, CRS, CRS + INT-777, and CRS + INT-777 + sdVx groups. Behavioral analyses and image quantification were performed by investigators blinded to group allocation where applicable.

### Chronic restraint stress model

Mice in the CRS groups were individually placed in ventilated 50-mL centrifuge tubes with openings at both ends. The restraint allowed slight head movement while restricting limb movement. Mice were subjected to restraint stress for 6 h per day for 4 consecutive weeks. During each restraint session, the animals were monitored regularly. After restraint, mice were returned to their home cages. Control mice were handled similarly but were not subjected to restraint stress.

### INT-777 preparation

INT-777 (MedChemExpress, USA) was dissolved in 2% carboxymethyl cellulose sodium. From day 15 of CRS exposure, mice in the CRS + INT-777 group received INT-777 by oral gavage at 30 mg/kg once daily for 14 consecutive days^29, 58, 59, 60^. Control and CRS mice received the corresponding vehicle. Body weight was recorded during the experimental period.

### Behavioral assays

#### Forced swimming test (FST)

Mice were placed individually in transparent glass cylinder (15 cm diameter × 30 cm height) filled with water to a depth of 15 cm at 25 ± 1 ℃. Behavior was recorded for 6 min, and immobility was analyzed during the last 5 min using behavioral tracking software^61^. Immobility was defined as the absence of active escape-related movements, with only the minimal movements necessary to keep the head above water.

### Tail suspension test (TST)

Mice were suspended individually by the tail using medical adhesive tape attached to the distal end of the tail and connected to a string fixed to a hook on a wooden suspension box, with no contact with nearby surfaces. Behavior was recorded for 6 min, and immobility was analyzed during the final 5 min with the EthoVision XT program (Noldus Information Technology BV, Wageningen, the Netherlands)^61^. Immobility was defined as the absence of active struggle, with only minimal movements related to respiration or postural maintenance.

### Open field test (OFT)

Each mouse was placed in the center of an open-field box (44 cm × 44 cm × 44 cm) and allowed to explore freely for 5.5 min^61^. A camera positioned directly above the arena was used to record the movement trajectory of each mouse and EthoVision XT software was used to quantify total distance traveled and time spent in the central zone. After each trial, urine and feces were removed, and the apparatus was thoroughly cleaned with 75% alcohol.

### Sucrose preference test (SPT)

Mice were provided with two bottles containing 200 mL of 1% sucrose solution and 200 mL of water, respectively, for sucrose adaptation over 3 consecutive days. The two bottles were placed in parallel on the feeding rack, and their positions were exchanged after 24 h to minimize side bias^61^. After the adaptation period, mice were deprived of water for 24 h, followed by simultaneous access to one bottle of 1% sucrose solution and one bottle of water. Fluid intake was recorded over the subsequent 24 h. Sucrose preference was calculated as the volume of sucrose solution consumed divided by the total fluid intake (sucrose solution + water) × 100%.

### Histological and Molecular Analyses Protein extraction and Western blotting

Prefrontal cortex, hippocampal, hypothalamic, and ileal tissue samples were homogenized on ice in RIPA with protease inhibitors and phenylmethylsulfonyl fluoride (PMSF). After centrifugation (4 ℃, 12000×g, 15 min), supernatant protein concentrations were determined by BCA assay. After protein extraction, samples were separated by sodium dodecyl sulfate-polyacrylamide gel electrophoresis (SDS-PAGE) and subsequently transferred onto polyvinylidene fluoride (PVDF) membranes (Millipore, USA). Blocking was performed using 5% non-fat milk at room temperature to prevent nonspecific binding. After blocking, the membranes were incubated with primary antibodies overnight at 4 ℃, followed by incubation with the corresponding HRP-conjugated secondary antibodies. Immunoreactive bands were detected using an enhanced chemiluminescence reagent and imaged with a Bio-Rad gel imaging system.

### Immunofluorescence assays

Mice were deeply anesthetized and transcardially perfused with PBS followed by 4% paraformaldehyde. Brains were post-fixed in 4% paraformaldehyde, dehydrated in sucrose, embedded in OCT, and sectioned coronally at 30 μm. Ileal tissues were fixed in 4% paraformaldehyde, paraffin-embedded, and sectioned at approximately 5 μm. After antigen retrieval, permeabilization, and blocking, sections were incubated overnight at 4 ℃ with the following primary antibodies: rabbit anti-Gpbar1 monoclonal antibody (Novus, Cat. No. NBP2-23669; 1:200), mouse anti-PGP9.5 monoclonal antibody (Proteintech, Cat. No. 66230-1-Ig; 1:200), anti-TUJ1 monoclonal antibody (Cell Signaling Technology, Cat. No. #5568; dilution [1:200]), anti-S100β monoclonal antibody (Proteintech, Cat. No. 66616-1-Ig; dilution [1:200]), rabbit anti-HK2 monoclonal antibody (Cell Signaling Technology, Cat. No. 55205; 1:200), rabbit anti-TOM20 monoclonal antibody (Cell Signaling Technology, Cat. No. 42406; 1:200), rabbit anti-GAD1 monoclonal antibody (Cell Signaling Technology, Cat. No. 41318; 1:50), rabbit anti-FosB monoclonal antibody (Cell Signaling Technology, Cat. No. 2251; 1:600), mouse anti-CaMKIIα monoclonal antibody (Cell Signaling Technology, Cat. No. 50049; 1:200), and rabbit anti-VGLUT1 monoclonal antibody (Novus, Cat. No. NBP2-59329; 1:200). Sections were then incubated with fluorescent secondary antibodies or processed using a multiplex fluorescence staining kit according to the manufacturer’s instructions. Nuclei were counterstained with DAPI. Images were acquired using fluorescence or confocal microscopy under identical settings across groups. Fluorescence intensity and colocalization were analyzed using Imaris and ImageJ.

### Untargeted metabolomics of ileal tissue

Terminal ileal tissues were collected, rinsed with precooled PBS, snap-frozen in liquid nitrogen, and sent to Majorbio Bio-Pharm Technology Co., Ltd. (Shanghai, China) for untargeted metabolomic profiling. Metabolites were extracted from ileal tissues and analyzed by ultra-high-performance liquid chromatography coupled with high-resolution mass spectrometry. Raw data were processed for peak detection, alignment, normalization, and metabolite annotation. PCA, OPLS-DA, volcano plot analysis, and KEGG pathway enrichment were used to identify differential metabolites and altered pathways.

### 16S rRNA Sequencing and Analysis

Ileal microbial samples were collected under sterile conditions and stored at -80° C until analysis. Total microbial DNA was extracted, and the bacterial 16S rRNA gene was amplified for high-throughput sequencing. Raw reads were quality-filtered, assembled, and cleared of chimeric sequences. Taxonomic annotation was performed against the SILVA database. Alpha diversity was assessed using the Chao, Shannon, and Simpson indices. Beta diversity was evaluated by principal coordinate analysis, and group differences were tested using Adonis analysis. Taxonomic composition at the genus and phylum levels was calculated and visualized as stacked bar plots.

### Targeted energy metabolomics of ileum and hypothalamus

Ileal and hypothalamic tissues were rapidly collected, frozen in liquid nitrogen, and sent to Majorbio Bio-Pharm Technology Co., Ltd. (Shanghai, China) for targeted energy metabolomics. Approximately 20 mg of tissue was extracted with 80% methanol, homogenized, sonicated, and centrifuged. Supernatants were analyzed using ultra-high-performance liquid chromatography-tandem mass spectrometry in multiple reaction monitoring mode. A targeted panel of energy-related metabolites was qualitatively and quantitatively analyzed.

### Lactate assay and ELISA

Hypothalamic tissues were homogenized on ice, and the supernatants were collected after centrifugation. Lactate content was measured using a lactate assay kit (Nanjing Jiancheng Bioengineering Institute, China) according to the manufacturer’s instructions. HK2 levels were measured using an enzyme-linked immunosorbent assay kit (Quanzhou Ruixin Biological Technology, China) according to the manufacturer’s protocol. Absorbance was measured using a microplate reader, and analytic concentrations were calculated from standard curves. When applicable, tissue levels were normalized to total protein concentration.

### Stereotaxic sparse-labeling viral injection and dendritic spine analysis

Mice were anesthetized and placed in a stereotaxic frame. CaMKIIα-FCSSP-EYFP-5E4 virus ( ≥ 2.00 × 10^12^ v.g./mL) was bilaterally injected into the hypothalamic region at AP −0.73 mm, ML ± 0.25 mm, and DV −4.75 mm^62^, with an injection volume of 300 nL per side at 60 nL/min. After injection, the capillary was kept in place briefly to minimize viral backflow, and mice were allowed to recover before subsequent experimental procedures.

### PRV-mediated retrograde transsynaptic tracing from the ileal wall

Mice were anesthetized and subjected to aseptic laparotomy. The terminal ileum was exposed under a stereomicroscope. PRV-CAG-EGFP (5 μ L; 2.00 × 10^9^ PFU/mL; BrainVTA, Wuhan, China) was injected into the ileal wall at multiple sites using a pulled glass capillary. The capillary was kept in place briefly to minimize viral leakage. The ileum was rinsed with sterile saline and returned to the abdominal cavity, and the abdominal wall was sutured in layers. Five days after viral inoculation, mice were deeply anesthetized and perfused. The ileum, nodose ganglion, and brain were collected, sectioned, and examined for EGFP expression signals. The NTS and PVN were identified according to the mouse brain atlas using the following coordinates: NTS, AP−7.4 mm, ML±0.5mm; DV-4.5mm; PVN, AP-0.73mm; ML±0.25mm; DV-4.75 mm^62^.

### Chemogenetic inhibition of PVN GABAergic neurons

For chemogenetic inhibition of PVN GABAergic neurons, we bilaterally injected pAAV-GAD67-hM4D(Gi)-mCherry into the PVN of mice in the CRS + INT-777 + hM4D(Gi) group and injected pAAV-GAD67-mCherry into the PVN of mice in the corresponding control groups. The PVN injection coordinates were AP −0.73 mm, ML ± 0.25 mm, and DV −4.75 mm. Before behavioral testing, we administered deschloroclozapine (DCZ; MCE, USA) intraperitoneally at 0.1 mg/kg to inhibit hM4D(Gi)-expressing PVN GABAergic neurons. We performed behavioral tests within the effective chemogenetic inhibition window. After behavioral experiments, we verified viral expression in the PVN and excluded animals with misplaced viral expression or insufficient fluorescence in the target region from the final analysis.

### Subdiaphragmatic vagotomy

Subdiaphragmatic vagotomy was performed to assess the involvement of vagal signaling in the behavioral effects of INT-777. One day before surgery, mice were given liquid food to reduce gastric contents. Under isoflurane anesthesia, the abdominal cavity was opened under aseptic conditions^63, 64, 65^. The stomach and lower esophagus were exposed, and the subdiaphragmatic vagal branch at the esophageal base was carefully transected using microsurgical scissors. The stomach was returned to the abdominal cavity, and the abdominal wall and skin were sutured in layers. Mice were allowed to recover for 1 week before CRS modeling, INT-777 administration, and behavioral testing.

### Statistical analysis

Unless otherwise specified, all data for bar charts are presented as mean ± standard error of the mean (SEM). Statistical analyses were performed using SPSS 26.0 and GraphPad Prism 9. Normality was assessed before statistical testing. For comparisons between two groups, unpaired two-tailed Student’s *t* test was used for normally distributed data, and the Mann–Whitney U test was used for non-normally distributed data. For comparisons among three or more independent groups, one-way ANOVA followed by Tukey’s post hoc test was used. Body weight changes over time were analyzed using two-way repeated-measures ANOVA followed by Bonferroni’s post hoc test when appropriate. For metabolomic data, PCA and OPLS-DA were used to evaluate group separation, and 200-permutation tests were performed to assess model robustness. Differential metabolites were selected according to VIP > 1 and *P*<0.05 unless otherwise specified. KEGG pathway enrichment analysis was performed for differential metabolites. Fluorescence intensity and colocalization were quantified using Imaris and ImageJ. Manders’ colocalization coefficients were calculated to quantify the degree of overlap between fluorescence signals. *P*<0.05 was considered statistically significant.

## Acknowledgements

This work was supported by the National Natural Science Youth Fund Project (82001686), National Natural Science Regional Foundation Project (82160273, 82160222), Basic Research Project of Guizhou Science and Technology Plan (Qiankehe Foundation ZK (2022) General 261), Guizhou High-Level Innovative Talent Program (Thousand Talent Tier, gzwjrs-021), Guizhou Provincial People’s Hospital National Science Foundation (GPPH-NSFC-2021-14), the Guizhou Provincial Clinical Medical Research Center Construction Project - Neurological Disease Research (No: LCZX[2025]003), the “ Summit Plan ” Project for the Construction of Clinical Key Specialties by Guizhou Provincial Health Commission (GZWJWDF2025003), and the Key Advantageous Discipline Construction Project of Guizhou Provincial Health Commission in 2023 (2023) China. Shan Guan acknowledges support for the research described in this study from the National Natural Science Foundation of China (NSFC, Grant No. 32370993), the National Science and Technology Major Project (Grant No. 2025ZD01903302, 2021YFC2302500 and 2024YFC2310804) by the Ministry of Science and Technology of China, and the Natural Science Foundation of Chongqing (CSTB2025NSCQ-GPX0624).

## Data availability

All relevant data supporting the key findings of this study are available within the article and its Supplementary files.

## Author contributions

T.T., L.Z. and S.G. conceived and designed the study. L.Z., Y.S., and M.Y. performed the behavioral tests, image acquisition, mouse experiments and data analysis. S.Y., J.W., D.G. performed stereotaxic injections. Z.C. provided experimental platform support. L.W., M.W., G.X., and T.T. acquired funding. Y.S. and M.Y. wrote the initial draft of the manuscript. S.G. and T.T. revised the manuscript and supervised the study. All authors read and approved the final manuscript.

