## Supplementary Information for "Ileal Gpbar1 gates depressive phenotypes via gut to brain “vagal-NTS-PVN” GABAergic axis"

**Contents**

Supplementary Table 1
**List of abbreviations**

Supplementary Figure 1-5

**List of abbreviations**

| **Abbreviation** | **Full name** |
| --- | --- |
| CRS | Chronic restraint stress |
| Gpbar1 | G protein-coupled bile acid receptor 1 |
| TGR5 | Takeda G protein-coupled receptor 5 |
| INT-777 | 6α-Ethyl-23(S)-methylcholic acid |
| NTS | Nucleus tractus solitarius |
| PVN | Paraventricular nucleus of the hypothalamus |
| GABA | γ-Aminobutyric acid |
| GAD1 | Glutamate decarboxylase 1 |
| HK2 | Hexokinase 2 |
| TOM20 | Translocase of outer mitochondrial membrane 20 |
| HIF-1α | Hypoxia-inducible factor 1α |
| mTOR | Mechanistic target of rapamycin |
| ATP | Adenosine triphosphate |
| PRV | Pseudorabies virus |
| EGFP | Enhanced green fluorescent protein |
| NG | Nodose ganglion |
| DREADD | Designer receptors exclusively activated by designer drugs |
| hM4D(Gi) | Gi-coupled inhibitory designer receptor hM4D |
| DCZ | Deschloroclozapine |
| sdVx | Subdiaphragmatic vagotomy |
| ENS | Enteric nervous system |
| MP | Myenteric plexus |
| SMP | Submucosal plexus |
| PGP9.5 | Protein gene product 9.5 |
| TUJ1 | Class III β-tubulin |
| S100β | S100 calcium-binding protein beta |
| CaMKIIα | Calcium/calmodulin-dependent protein kinase II alpha |
| VGLUT1 | Vesicular glutamate transporter 1 |
| PSD95 | Postsynaptic density protein 95 |
| OFT | Open field test |
| SPT | Sucrose preference test |
| FST | Forced swimming test |
| TST | Tail suspension test |
| 16S rRNA | 16S ribosomal RNA |
| PCoA | Principal coordinate analysis |
| KEGG | Kyoto Encyclopedia of Genes and Genomes |
| PCA | Principal component analysis |
| SEM | Standard error of the mean |
| SDS-PAGE | Sodium dodecyl sulfate-polyacrylamide gel electrophoresis |
| PVDF | Polyvinylidene fluoride |
| BCA | Bicinchoninic acid assay |
| PMSF | Phenylmethylsulfonyl fluoride |

**SUPPLEMENTARY FIGURES**

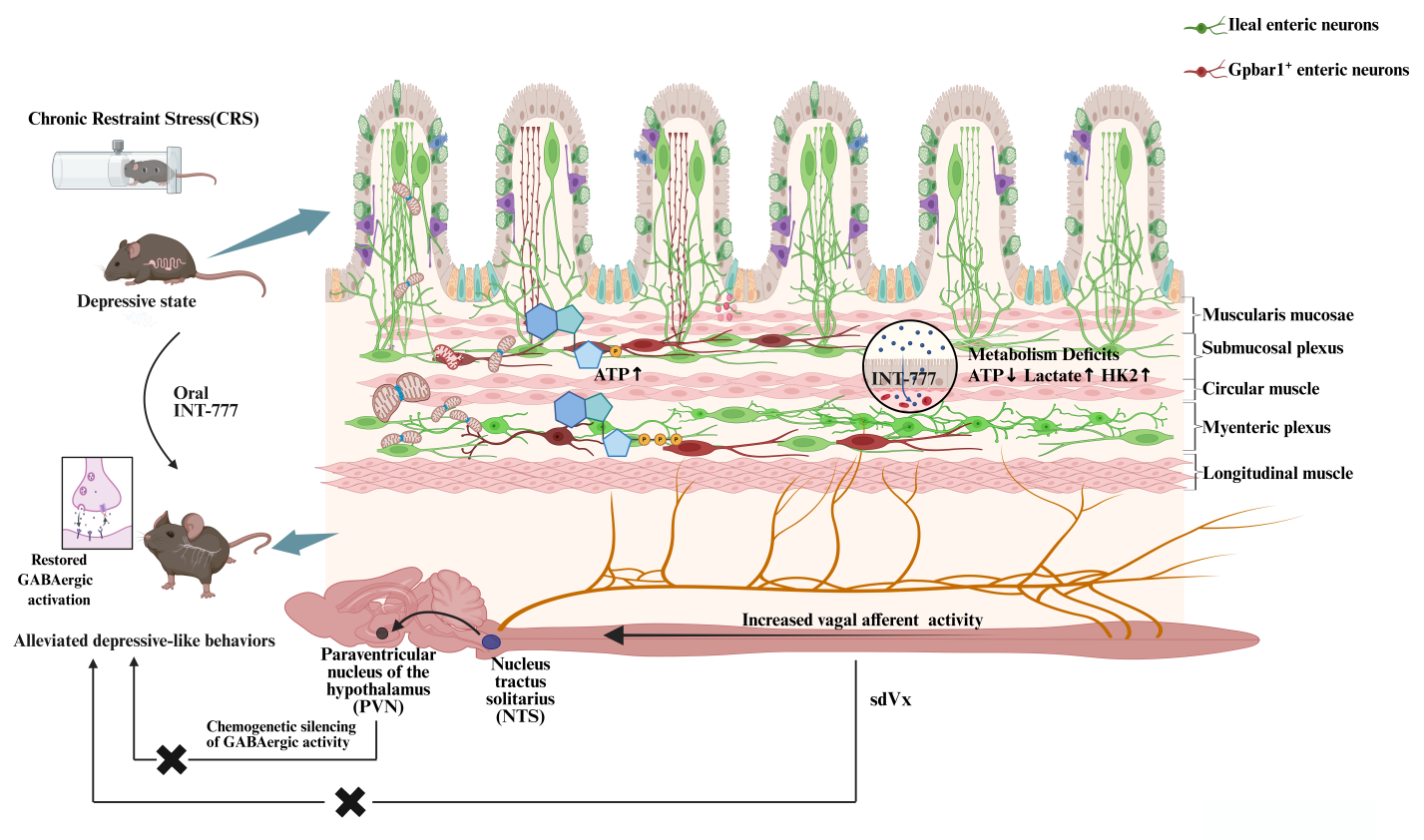

**Supplementary Fig. 1| Ileal enteric neuronal Gpbarl activation modulates gut-brain energy metabolism and ileal vagal afferent-NTS-PVNGABAergic signaling to alleviate depression-like behaviors.**

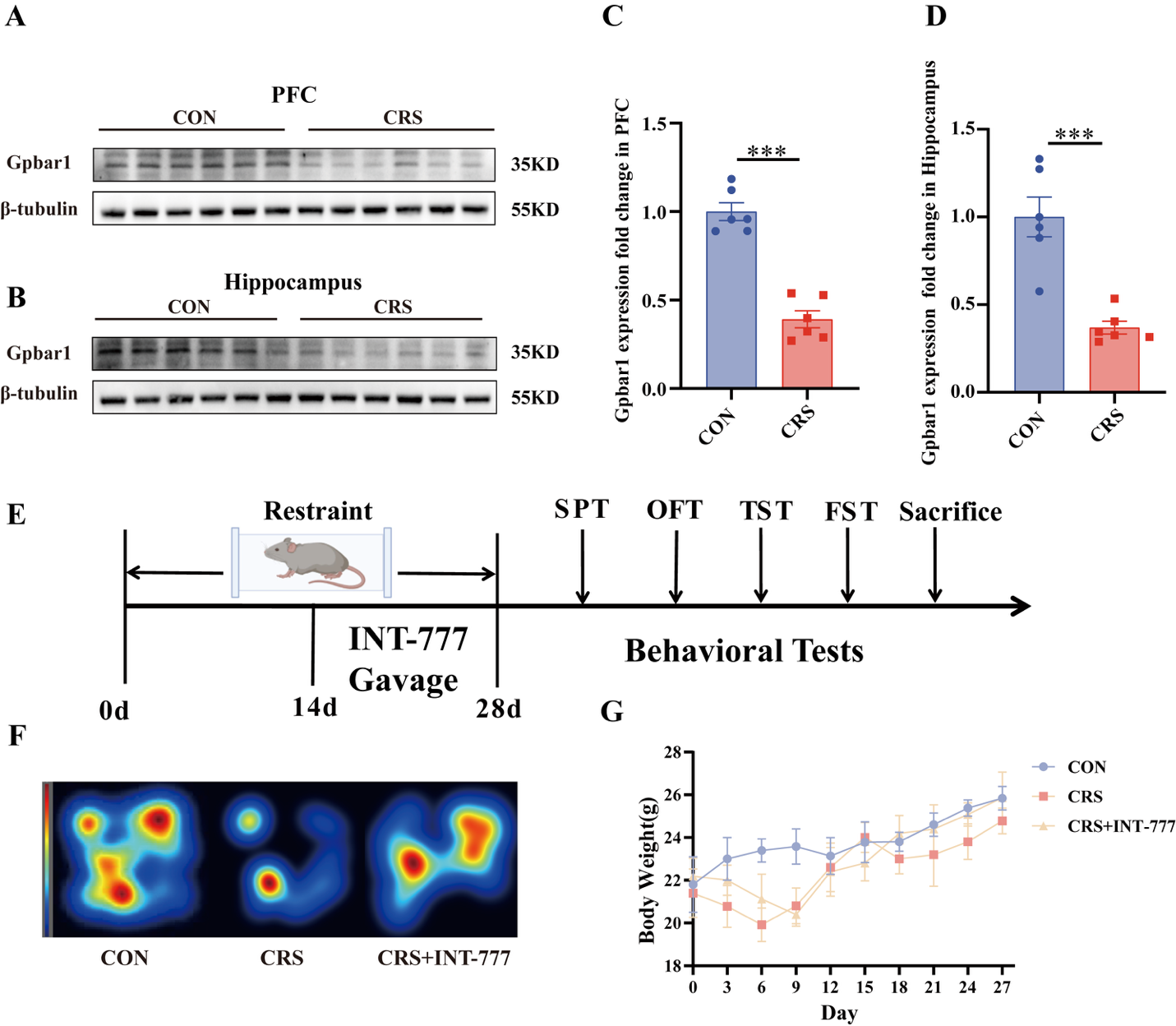

**Supplementary Fig. 2| CRS reduces Gpbar1 expression in the prefrontal cortex and hippocampus, with experimental timeline, movement heatmaps, and body weight changes after INT-777 intervention.** (A) Representative western blots of Gpbar1 expression in the prefrontal cortex of CON and CRS mice. (B) Representative western blots of Gpbar1 expression in the hippocampus of CON and CRS mice. (C) Quantification of Gpbar1 protein expression in the prefrontal cortex. (D) Quantification of Gpbar1 protein expression in the hippocampus. (E) Experimental timeline of CRS exposure, INT-777 administration by oral gavage, behavioral testing, and tissue collection. (F) Representative movement-density heatmaps showing center-zone exploration in the open field test across the CON, CRS, and CRS + INT-777 groups. (G) Body weight changes recorded every 3 days during the experimental period.Data are presented as mean ± SEM. ns, not significant; *P < 0.05, **P < 0.01, and ***P < 0.001.

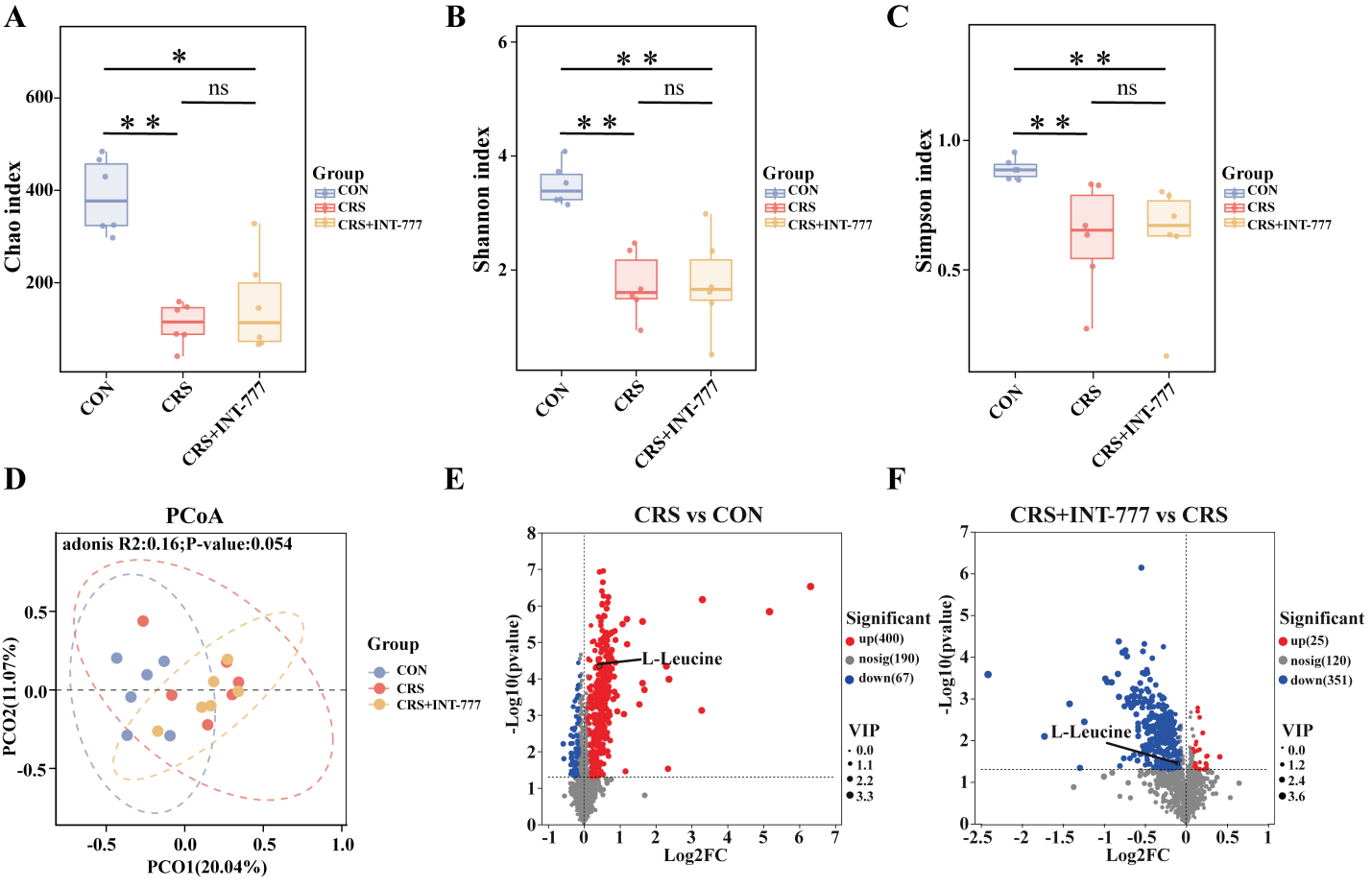

**Supplementary Fig. 3| INT-777 does not restore ileal microbiota diversity but modulates ileal metabolic alterations in CRS mice. (A)** Chao index of ileal microbial communities among the CON, CRS, and CRS + INT-777 groups. **(B)** Shannon index of ileal microbial communities among the three groups. **(C)** Simpson index of ileal microbial communities among the three groups. **(D)** Principal coordinate analysis showing the distribution of ileal microbial communities among the CON, CRS, and CRS + INT-777 groups. Adonis analysis showed a separation trend among groups without reaching statistical significance. **(E)** Volcano plot showing differentially abundant ileal metabolites between the CRS and CON groups, with L-leucine labeled. **(F)** Volcano plot showing differentially abundant ileal metabolites between the CRS + INT-777 and CRS groups, with L-leucine labeled. Data are presented as mean ± SEM. ns, not significant; *P < 0.05, **P < 0.01, and ***P < 0.001.

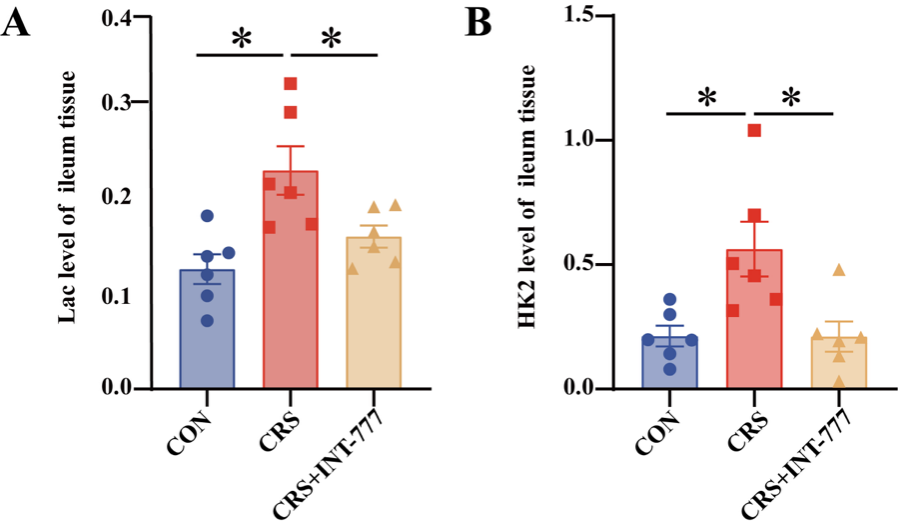

**Supplementary Fig. 4| INT-777 ameliorates CRS-induced glycolytic alterations in ileal tissue.** (A) Lactate levels in ileal tissues among the CON, CRS, and CRS + INT-777 groups. (B) HK2 levels in ileal tissues among the CON, CRS, and CRS + INT-777 groups. Data are presented as mean ± SEM. ns, not significant; *P < 0.05, **P < 0.01, and ***P < 0.001.

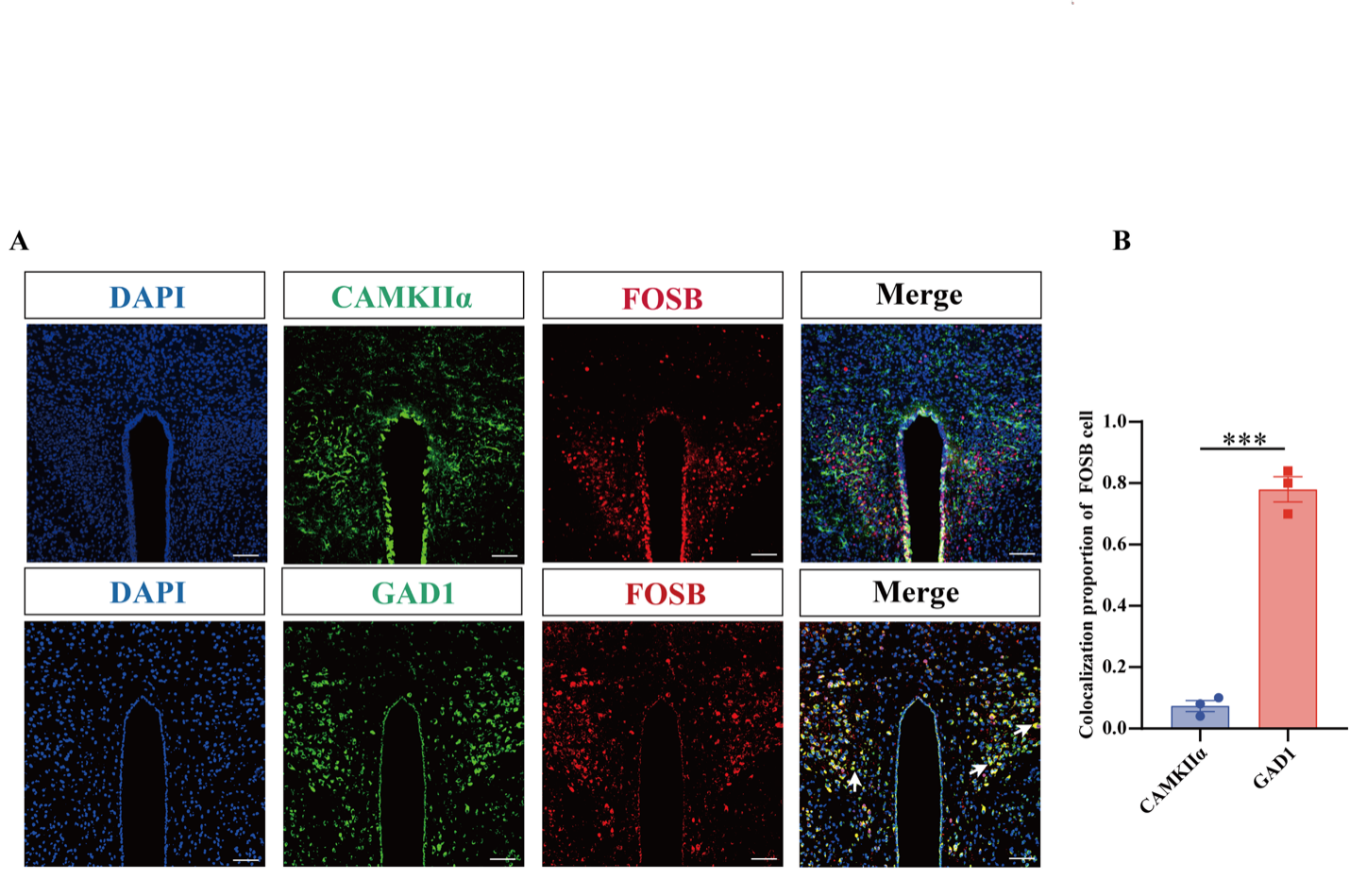
 **Supplementary Fig. 5| FosB-positive signals in the PVN are preferentially associated with GAD1-positive neurons.** (**(A)** Representative immunofluorescence images showing FosB co-staining with CaMKIIα or GAD1 in the PVN. DAPI, blue; CaMKIIα or GAD1, green; FosB, red. White arrows indicate representative co-localized signals. **(B)** Quantification of the colocalization proportion of FosB-positive signals with CaMKIIα-positive or GAD1-positive neurons in the PVN. Data are presented as mean ± SEM. ns, not significant; *P < 0.05, **P < 0.01, and ***P < 0.001.
